# Kaposi’s sarcoma-associated herpesvirus fine-tunes the temporal expression of late genes by manipulating a host RNA quality control pathway

**DOI:** 10.1101/2020.02.18.955526

**Authors:** Julio C. Ruiz, Anne Devlin, Jiwoong Kim, Nicholas K. Conrad

## Abstract

Kaposi’s sarcoma-associated herpesvirus (KSHV) is a human oncogenic nuclear DNA virus that expresses its genes using the host cell transcription and RNA processing machinery. As a result, KSHV transcripts are subject to degradation by at least two host-mediated nuclear RNA decay pathways, PABPN1 and PAPα/γ-mediated RNA decay (PPD) and an ARS2-dependent decay pathway. Here, we present global analyses of viral transcript levels to further understand the roles of these decay pathways in KSHV gene expression. Consistent with our recent report that the KSHV ORF57 protein increases viral transcript stability by impeding ARS2-dependent decay, ARS2 knockdown has little effect on viral gene expression 24 hours after lytic reactivation of wild-type virus. In contrast, inactivation of PPD results in premature accumulation of late transcripts. The up-regulation of late transcripts does not require the primary late gene-specific viral transactivation factor, suggesting that cryptic transcription produces the transcripts that then succumb to PPD. Remarkably, PPD inactivation has no effect on late transcripts at their proper time of expression. We show that this time-dependent PPD evasion by late transcripts requires the host factor NRDE2, which has previously been reported to protect cellular RNAs by sequestering decay factors. From these studies, we conclude that KSHV uses PPD to fine-tune the temporal expression of its genes by preventing their premature accumulation.

**Importance:** Kaposi’s sarcoma-associated herpesvirus (KSHV) is an oncogenic gammaherpesvirus that causes Kaposi’s sarcoma and other lymphoproliferative disorders. Nuclear expression of KSHV genes results in exposure to at least two host-mediated nuclear RNA decay pathways, PABPN1 and PAPα/γ-mediated RNA decay (PPD) and an ARS2-mediated decay pathway. Perhaps unsurprisingly, we previously found that KSHV uses specific mechanisms to protect its transcripts from ARS2-mediated decay. In contrast, here we show that PPD is required to dampen the expression of viral late transcripts that are prematurely transcribed, presumably due to cryptic transcription early in infection. At the proper time for their expression, KSHV late transcripts evade PPD through the activity of the host factor NRDE2. We conclude that KSHV fine-tunes the temporal expression of its genes by modulating PPD activity. Thus, the virus both protects from and exploits the host nuclear RNA decay machinery for proper expression of its genes.

## Introduction

Kaposi’s sarcoma-associated herpesvirus (KSHV; also known as human herpesvirus 8; HHV-8), is an enveloped, double-stranded DNA virus. It is the etiological agent of Kaposi’s sarcoma and of the lymphoproliferative disorders primary effusion lymphoma (PEL) and multicentric Castleman’s disease (MCD) (1–4). Like other herpesvirus, the KSHV life cycle consists of a latent phase and a lytic phase. During latency, the viral genome resides in the host nucleus as a non-integrated, circular episome, and no virions are produced. Upon reactivation, the virus undergoes a well-regulated cascade of gene expression initiated by the viral transactivator ORF50 (Rta) that ultimately results in the production of infectious virus (5–7). KSHV transcription and genome replication occur in the nucleus where the virus takes control of the host machinery needed for these processes. Consequently, similar to host RNAs, viral transcripts are subject to host-mediated RNA quality control (QC) pathways (8–12).

RNA QC pathways play an essential role during RNA biogenesis (8–12). In addition to eliminating misprocessed transcripts, RNA QC pathways prevent the accumulation of unstable, non-coding RNAs such as <u>prom</u>oter-u<u>p</u>stream <u>t</u>ranscripts (PROMPTs, also called uaRNAs) (13–15). PROMPTs are polyadenylated, non-coding RNAs with no or few introns that are transcribed from bidirectional promoters antisense to protein-coding genes (16–18). Accumulation of PROMPTs has deleterious effects for the cells as they compete with coding transcripts for the translational machinery (15). In eukaryotes, at least two nuclear RNA decay pathways prevent the accumulation of PROMPTs (13–15, 19, 20). Primarily, they are degraded through the CBCN complex that is recruited to the RNA via its 5’ cap. CBCN consists of the cap-binding complex (CBC), the ARS2 protein, and the nuclear exosome targeting (NEXT) complex (13, 21). The NEXT complex subunit MTR4 recruits the RNA exosome to degrade the transcript (13, 22).

PROMPTs and other long polyadenylated nuclear RNAs are also degraded by the PABPN1 and PAPα/γ-mediated RNA decay (PPD) pathway in which decay factors are recruited through 3’ poly(A) tail (14, 15, 19, 20, 23, 24). In this pathway, the nuclear poly (A)-binding protein (PABPN1) promotes poly(A) tail extension of target transcripts by stimulating the function of the poly (A) polymerases (PAPα or PAPγ; abbreviated PAPα/γ). The targeted RNAs are subsequently degraded by the nuclear RNA exosome. Recruitment of the exosome to polyadenylated RNAs is mediated by the zinc finger protein ZFC3H1, which links the exosome cofactor MTR4 to PABPN1. This link was coined the poly(A) tail exosome targeting (PAXT) connection (also called the polysome protector complex, PPC) (15, 20). Overall, both PPD and the CBCN complex survey the integrity of RNAs to eliminate transcriptional noise, misprocessed and other potentially detrimental RNAs.

ARS2 directly interacts with the CBC to form a hub that allows the assembly of mutually exclusive complexes that dictate the fate of a transcript (13, 21, 24, 25). For instance, ARS2 interacts with PHAX or ALYREF to promote the nuclear export of properly processed snRNAs and mRNAs, respectively (21, 24–26). Alternatively, ARS2 recruits NEXT or PAXT to target RNAs for exosome-mediated degradation (13, 20, 27). In some cases, ARS2 targets transcripts for degradation independently of NEXT or PPD/PAXT (28, 29). These complex interconnections challenge efforts to uncover the degree of independence or redundancy between nuclear RNA decay pathways. Nonetheless, ARS2 clearly plays a central role in promoting the decay of a number of nuclear transcripts.

Like their host counterparts, KSHV mRNAs are capped and polyadenylated, but most KSHV genes are short and intronless (30, 31). Consequently, cellular RNA QC pathways may degrade KSHV RNAs due to their similarity to PROMPTs. The essential multifunctional KSHV ORF57 protein promotes viral transcript accumulation by increasing nuclear RNA stability (32–45). We recently reported that viral transcripts are subject to degradation by both PPD and an ARS2-dependent but NEXT-independent decay pathway upon lytic reactivation of virus lacking ORF57 (29). Using pulse-chase assays with an unstable form of the KSHV nuclear non-coding PAN RNA (PAN∆ENE), we further showed that ORF57 preferentially protects viral transcripts from the ARS2-dependent decay pathway (29). Interestingly, although viral transcripts succumb to PPD, ORF57 protection of PAN∆ENE from PPD was modest, suggesting the possibility that a subset of viral transcripts undergo PPD-dependent degradation in the presence of ORF57 (29). However, the role of PPD during viral infection with ORF57 expressing virus has not been fully explored.

Here, we used RNAi to inactivate PPD and/or ARS2-dependent decay and monitored their contributions to KSHV gene expression in the presence of ORF57 by RNA-seq at 24 hours after lytic reactivation. ARS2 depletion resulted in few changes in viral gene expression. However, PPD inactivation resulted in increased expression levels of several viral genes. Interestingly, the most upregulated transcripts were late transcripts that are otherwise expressed at ~48 hours after lytic reactivation. Our data suggest that PPD prevents the premature accumulation of late transcripts which presumably arise as a consequence of cryptic transcription. Notably, at their proper time of expression, PPD inactivation has no effect on viral late transcripts, and the host factor NRDE2 is needed for evasion of PPD. We conclude that KSHV exploits PPD to fine-tune the temporal expression of viral genes by dampening steady-state levels of prematurely transcribed late genes.

## Results

### PPD inactivation results in aberrant temporal expression of KSHV late genes

Our previous studies showed that viral RNAs were subject to both PPD and ARS2-mediated decay in the absence of ORF57 (29). ORF57 more potently protected viral transcripts from ARS2-mediated decay than from PPD, suggesting that viral transcripts succumb to PPD even in the presence of ORF57. To further characterize the role of these decay pathways during KSHV infection, we performed an RNA-seq experiment to monitor the levels of viral transcripts after siRNA depletion of ARS2, the PPD component PAPα/γ, or both simultaneously (dKD) in iSLK cells latently infected with the KSHV infectious clone BAC16 (iSLK WT) (46, 47)(Fig 1A). Efficiency of knockdown was validated by western blot, qRT-PCR and/or loss of function assays (Fig 1B). Lytic reactivation was induced using doxycycline (dox) to promote expression of the dox-inducible RTA integrated into the iSLK host cell chromosomes and by the histone deacetylase inhibitor sodium butyrate (NaB). We prepared libraries from RNA harvested 24 hours post induction (hpi), and the samples were subjected to high-throughput sequencing (Fig 1A). Expression of several KSHV genes significantly increased in samples depleted of PAPα/γ and in the dKD compared to samples treated with a control siRNA. However, we observed minimal alterations in gene expression in the ARS2 knockdown samples, consistent with the idea that ARS2-mediated decay is inhibited by ORF57 (Fig 1C and Table S1). Surprisingly, the most upregulated genes (> 4-fold change) upon PAPα/γ depletion and in the dKD were late genes, 76% and 70% respectively (Fig 1D). Typically, these KSHV late genes are not expressed at ~24 hpi in these cells. These data suggest that KSHV exploits PAPα/γ to temporally control the expression of late genes.

**Fig 1.**
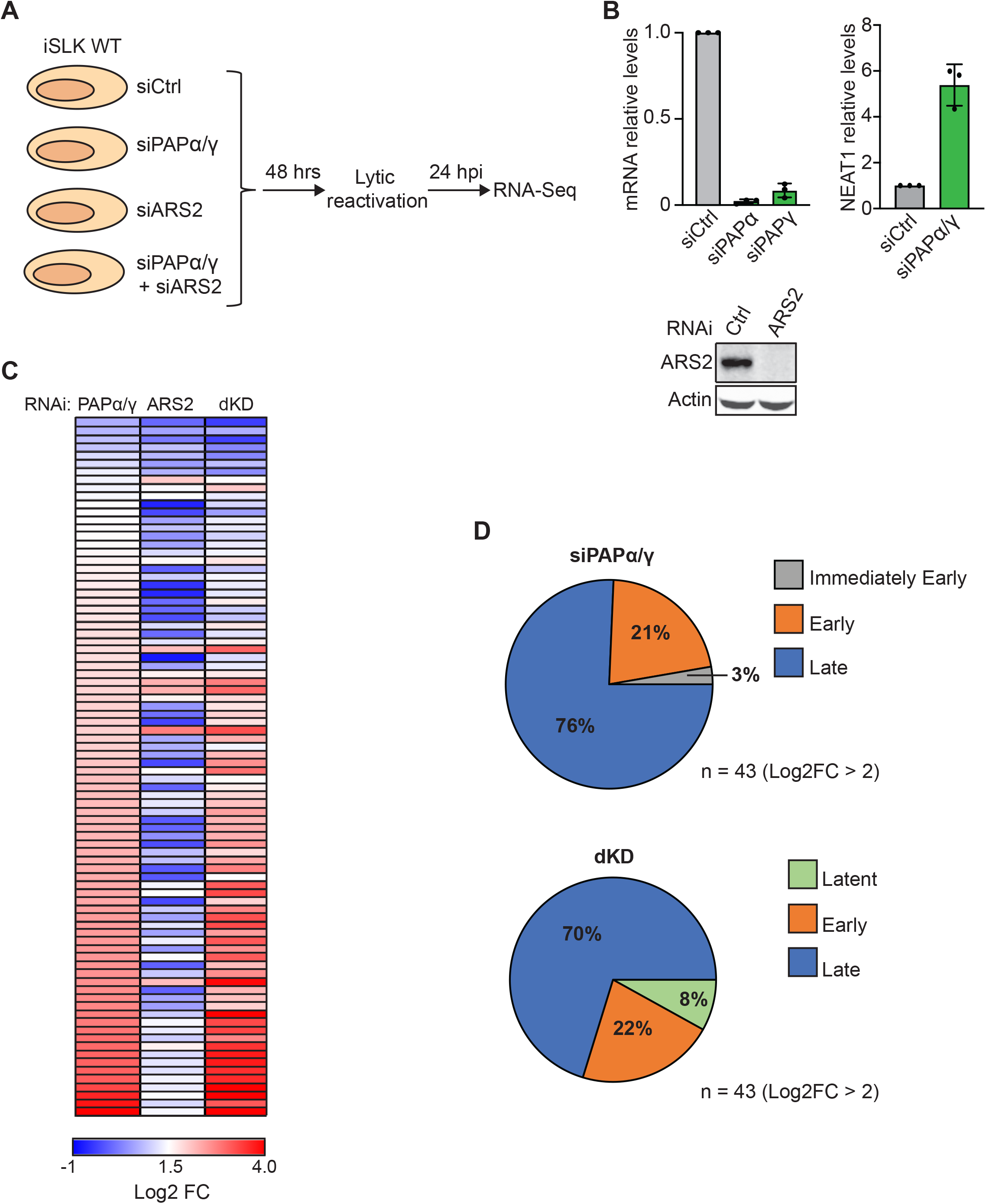
PPD inactivation affects the temporal expression of KSHV late genes. (A) Diagram of the RNA-seq experiment. iSLK WT cells were transfected with a non-targeting control siRNA or a two-siRNA pool targeting PAPα/γ, ARS2, or both PAPα/γ and ARS2 combined (dKD). Total RNA was harvested three days after siRNA transfection and 24 hours after lytic reactivation. Stranded mRNA-seq libraries were prepared and sequenced. (B) Efficiency of knockdown of PAPα, PAPγ, and ARS2 in iSLK WT cells. Due to the lack of robust antibodies, PAPα and PAPγ knockdown efficiency was determined by qRT-PCR. Bar graphs show PAPα and PAPγ mRNA levels in iSLK WT cells treated with siRNAs targeting PAPα and PAPγ. Because RNA knockdown does not necessarily correlate with protein loss, we assayed for loss of functional activity. To do so, we measured the RNA levels of a known PPD target, NEAT1, under the same conditions used for RNA-seq. Bar graph shows NEAT1 levels determined by qRT-PCR in iSLK WT cells depleted of PAPα/γ. Values are displayed relative to siCtrl after normalization to the 18S rRNA level. All values are averages, and the error bars are standard deviations (n = 3). ARS2 knockdown efficiency was determined by quantitative western blot. Actin serves as loading control. (C) Heatmap showing the log2 fold change (FC) relative to siCtrl for all KSHV genes in samples depleted of PAPα/γ, ARS2 or dKD. Genes are arranged in increasing log2 FC order based on PAPα/γ, where red represents maximum fold change and blue represents Log2 FC <1.5. (D) Pie charts showing the distribution of upregulated (>4-fold) KSHV genes in PAPα/γ (top) and dKD (bottom) according to their phase of expression (30).

### PPD/PAXT targets KSHV late genes for degradation

Our RNA-seq data show that PAPα/γ depletion results in increased expression of late genes at 24 hpi and suggest that PPD controls the expression late genes that prematurely arise due to cryptic transcription. To validate these observations and extend the findings to additional PPD factors, we focused our attention on three KSHV late genes, ORF52, ORF75, and K8.1. These genes were selected because they have different structural features. ORF52 is a short transcript (395 base pairs (bp)) while ORF75 is a long transcript (3890 bp). Both ORF52 and ORF75 are intronless, but K8.1 contains one intron. In spite of these structural differences, the expression of each gene increased upon PAPα/γ depletion at 24 hpi (Fig 2B, D and F). None were affected by ARS2 depletion alone. Similar results were obtained when the mRNA levels of these genes were monitored by qRT-PCR (Fig 2C, E and G). In principle, PAPα/γ knockdown may affect gene expression independent of PPD due to its function in 3’-end formation. To confirm the role of PPD, we tested whether depletion of PPD factors other than PAPα/γ caused a similar phenotype. We depleted cells of the unique PPD/PAXT factor ZFC3H1 or the exosome co-factor MTR4. Efficiency of knockdown was monitored by western blot, qRT-PCR and loss of function assays (Fig 2A). Consistent with PPD inactivation, depletion of ZFC3H1 or MTR4 also increased ORF75, ORF52 and K8.1 levels (Fig 2C, E and G). These data support the conclusion that PPD/PAXT suppresses KSHV late gene expression during the early stages of the virus lytic phase.

**Fig 2.**
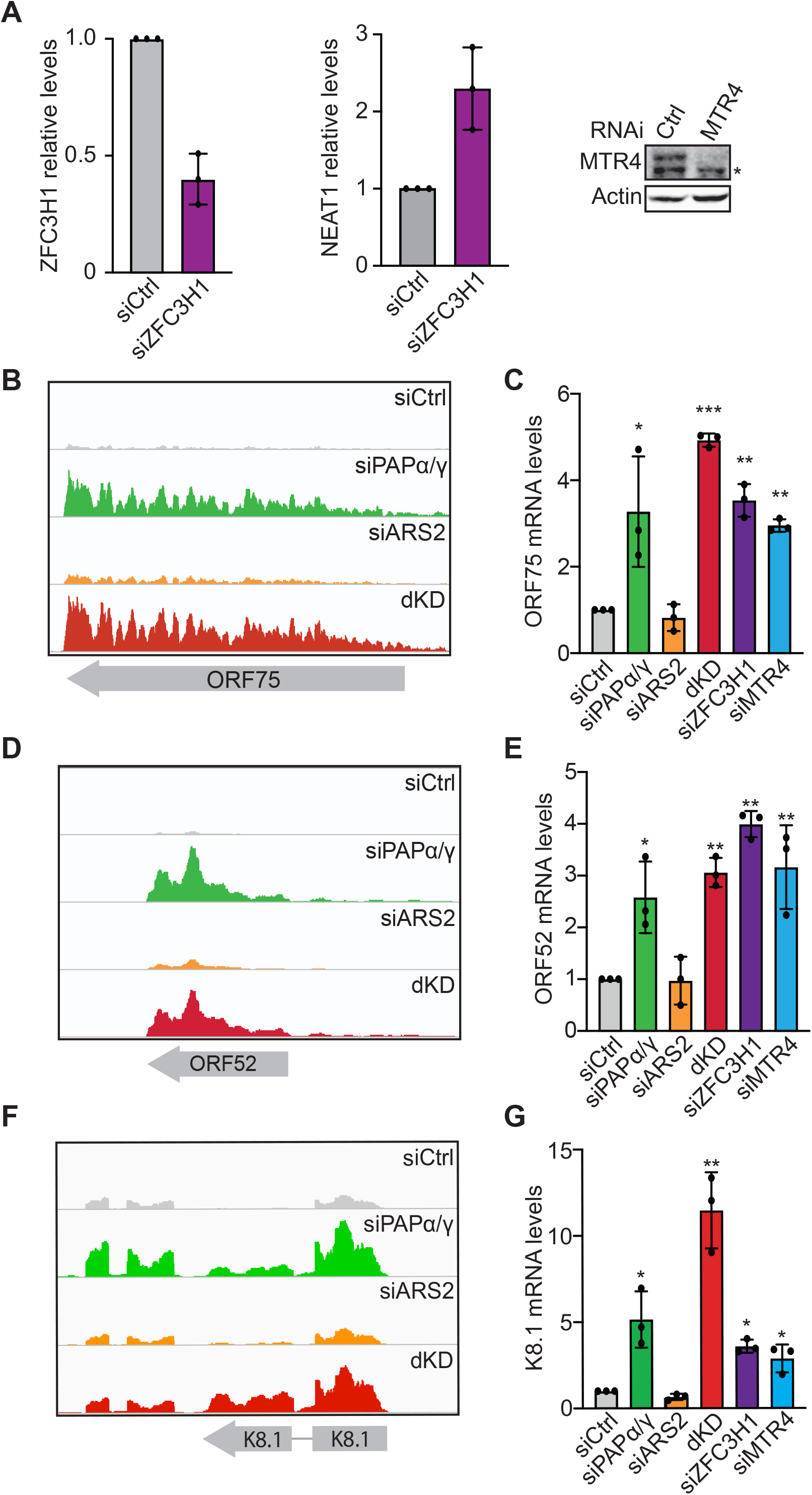
KSHV late transcripts are degraded by PPD/PAXT. (A) Efficiency of knockdown of ZFC3H1 and MTR4 in iSLK WT cells. Bar graphs show ZFC3H1 and NEAT1 mRNA levels in iSLK WT cells treated with ZFC3H1 siRNAs. Increased levels of NEAT1 indicate that PPD was effectively inactivated upon ZFC3H1 depletion. MTR4 knockdown efficiency was determined by quantitative western blot. Actin serves as loading control. (*) nonspecific band. (B, D and F) Integrative genome viewer (IGV) browser screenshots showing ORF75 (B), ORF52 (D) and K8.1 (F) reads in samples depleted of PAPα/γ, ARS2 or dKD. Each condition is depicted at the same scale. (C, E and G) Bar graphs showing relative ORF75 (C), ORF52 (E) and K8.1 (G) mRNA levels in iSLK WT cells depleted of PAPα/γ (green), ARS2 (orange), dKD (red), ZFC3H1 (purple) and MTR4 (light blue). Total RNA was harvested 24 hpi and analyzed by qRT-PCR. Values are displayed relative to siCtrl after normalization to the 18S rRNA level. All values are averages, and the error bars are standard deviations (n = 3). *P* values were determined by two-tailed unpaired Student’s *t* test: * < 0.05; ** < 0.01; *** < 0.001.

### PPD regulates virus late gene expression independently of the viral transactivation factor ORF24

To elucidate the mechanism of late gene expression upregulation in the context of PPD inactivation, we first focused our attention on the requirements for late gene expression. In KSHV, transcription of late genes takes place after initiation of viral DNA replication, and their expression requires the action of several viral transactivation factors (vTF) (48–53). The viral TATA box-binding protein homolog, ORF24, is a KSHV transactivation factor that plays a critical role when the virus progresses from DNA replication to expression of late genes. ORF24 binds to late gene promoters and recruits RNA polymerase II (pol II) and other vTFs to the promoter region of late genes to induce their expression. Because ORF24 mRNA levels were increased in cells depleted of PAPα/γ and in the dKD at 24 hours post lytic reactivation (Fig 3A and 3B), it is possible that PPD inactivation results in ORF24 upregulation which drives the increased expression of late genes. In this case, the effects of PPD on other late transcripts would result from a secondary effect of ORF24 upregulation. To determine whether the increased expression of late genes at 24 hpi is due to an upregulation of ORF24 upon PPD inactivation, we used iSLK cells transfected with a bacmid encoding the viral genome in which ORF24 contains a point mutation, R328A, that renders it inactive (51). ORF24^R328A^ maintains the interaction with pol II but is unable to interact with other vTAs resulting in strongly impaired expression of late genes (Fig 3C). If the up-regulation of late genes by PPD inactivation is due to secondary effects of ORF24 upregulation, then the up-regulation will be abrogated in the mutant virus. In contrast to this prediction, depletion of PPD components PAPα/γ, MTR4 or ZFC3H1 resulted in increased mRNA levels of ORF52, ORF75 and K8.1 in ORF24^R328A^ reactivated cells (Fig 3D, E and F). Thus, the role of PPD in late gene expression at 24 hpi is independent of ORF24 transactivation. Moreover, these data are consistent with a proposed role for PPD in the posttranscriptional inhibition of premature late transcript accumulation.

**Fig 3.**
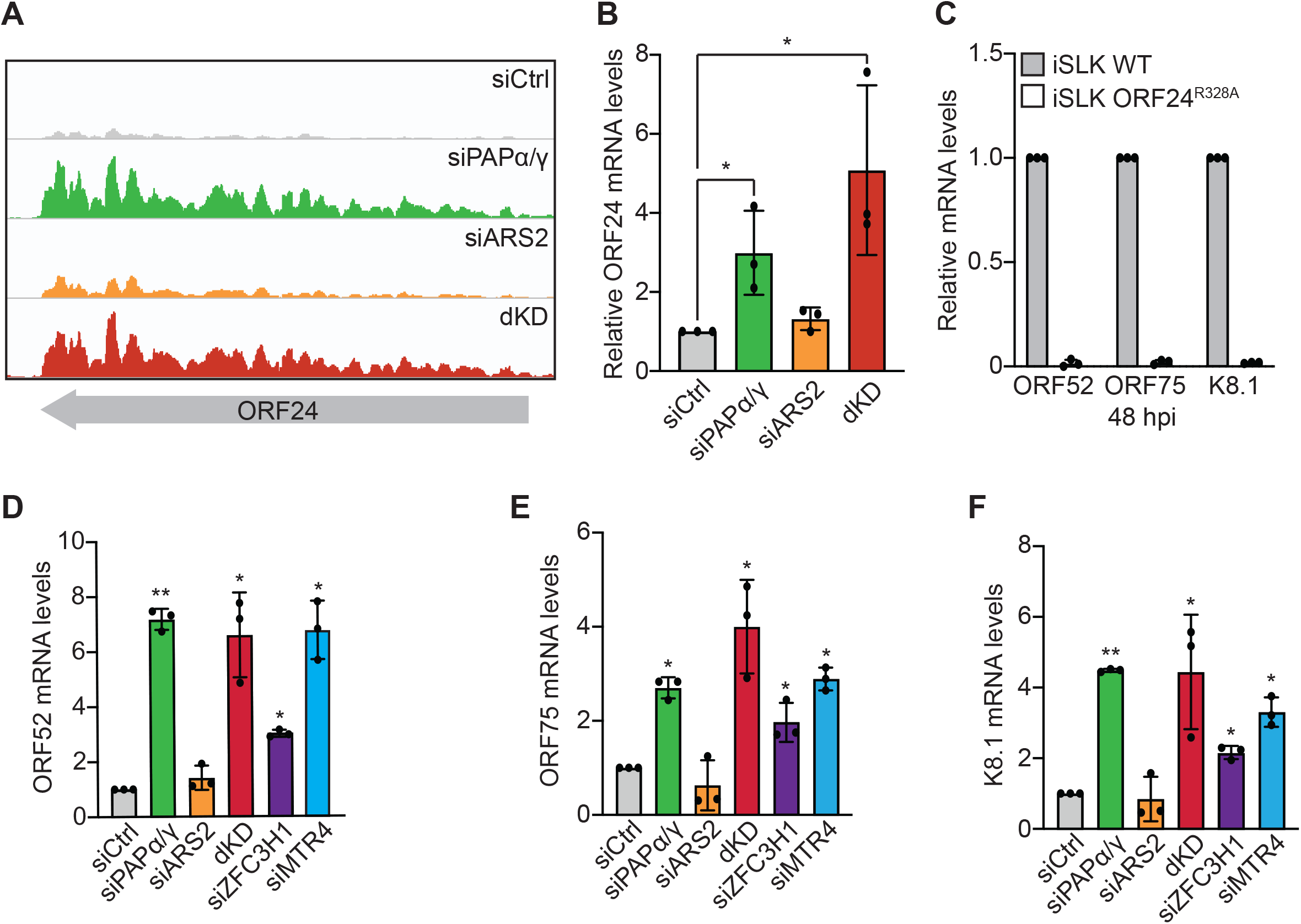
PPD upregulation of late genes is independent of ORF24. (A) IGV browser screenshot showing ORF24 reads in samples depleted of PAPα/γ, ARS2 or dKD. (B) Bar graphs showing relative ORF24 mRNA levels in iSLK cells depleted of PAPα/γ (green), ARS2 (orange) and dKD (red). (C) Bar graphs showing relative ORF52, ORF75 and K8.1 mRNA level in iSLK WT (gray) and iSLK ORF24^R328A^ (white) cells at 48 hpi. (D, E and F) Bar graphs showing relative ORF52 (D), ORF75 (E) and K8.1 (F) mRNA levels in iSLK ORF24^R328A^ cells depleted of PAPα/γ (green), ARS2 (orange), dKD (red), ZFC3H1 (purple) and MTR4 (light blue). Total RNA was harvested 24 hpi and analyzed by qRT-PCR. Values are displayed relative to siCtrl after normalization to the 18S rRNA level. All values are averages, and the error bars are standard deviations (n = 3). *P* values were determined by two-tailed unpaired Student’s *t* test: * < 0.05; ** < 0.01; *** < 0.001.

### PPD control of late transcripts is restricted to early phases of reactivation but does not affect genome replication or virus production in iSLK cells

KSHV late genes are expressed after the initiation of viral genome replication (48–52), which occurs prior to 48 hpi in our iSLK WT cells. As PPD inactivation increases late gene expression at 24 hpi, we tested whether the expression of late genes at 48 and 72 hpi is also affected by PPD inactivation. Interestingly, at 48- or 72-hpi, PAPα/γ depletion had no effect on the expression levels of the late genes tested (Fig 4A, B and C). These data suggest that at the proper time of expression, KSHV late transcripts are able to avoid PPD-mediated degradation. They further support the model that PPD dampens premature expression of KSHV late genes.

**Fig 4.**
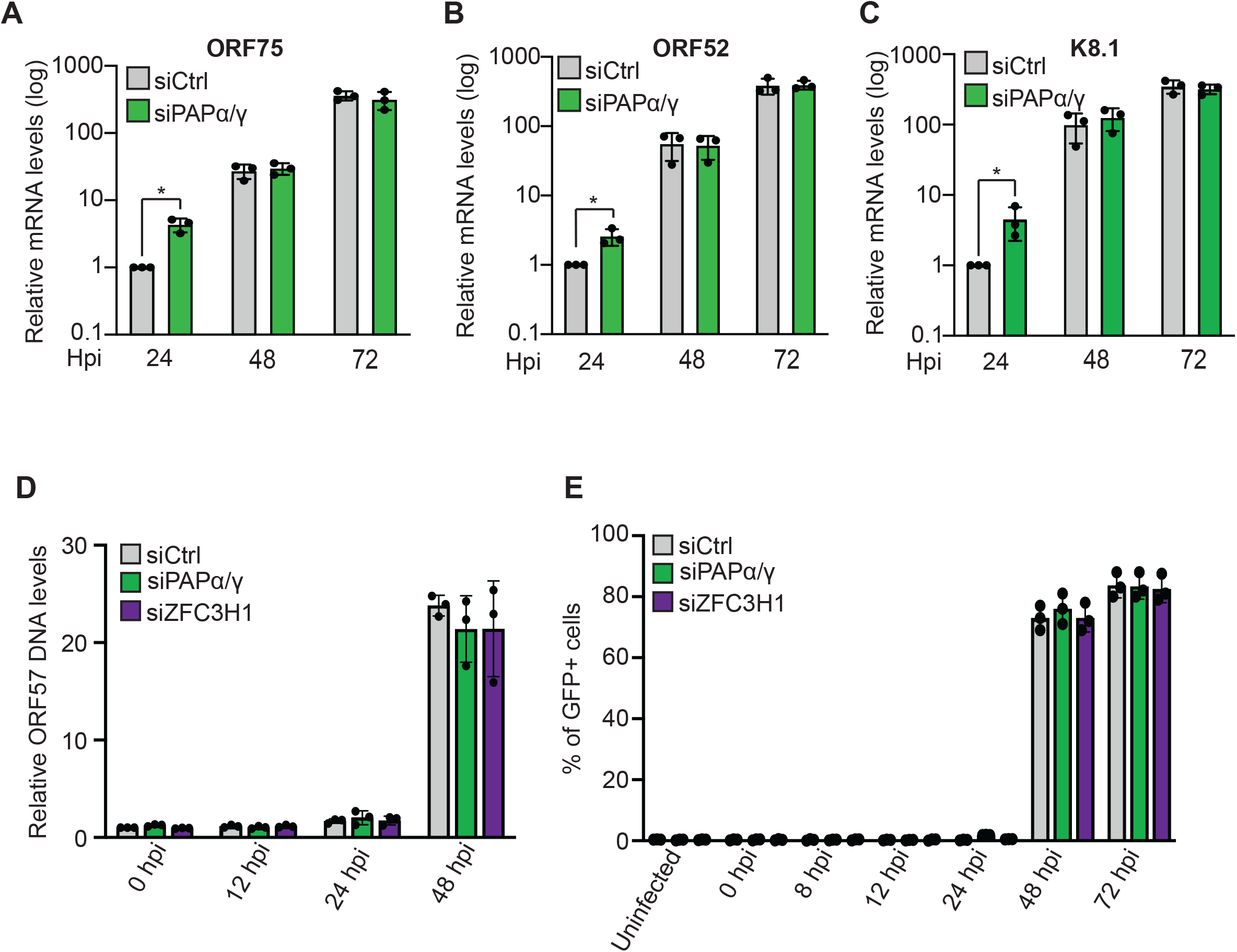
KSHV late transcripts evade PPD degradation at their proper time of expression. (A, B and C) Bar graphs showing relative ORF75 (A), ORF52 (B) and K8.1 (C) mRNA levels in iSLK WT cells treated with siRNAs targeting PAPα/γ (green) or a control siRNA (gray). Total RNA was harvested at 24, 48 and 72 hpi. Values were calculated relative to siCtrl at 24 hpi (gray) and normalized to the 18S rRNA level. Note that the data are plotted on a log scale due to the strong up-regulation of late genes after 48 hpi. (D) Bar graphs showing relative ORF57 DNA levels in iSLK WT cells depleted of PAPα/γ (green) or ZFC3H1 (purple). DNA was harvested at 0, 12, 24 and 48 hpi. Values were calculated relative to siCtrl (gray) at 0 hpi. (E) Bar graph of flow cytometry analysis showing percentage of GFP-positive HEK293 cells infected with supernatants collected from iSLK WT cells at 0, 8, 12, 24, 48 and 72 hpi. All values are averages, and the error bars are standard deviations (n = 3). *P* values were determined by Student’s *t* test: * < 0.05; ** < 0.01; *** < 0.0001.

Because of the aberrant timing of viral late gene expression, we investigated whether PPD inactivation affected virus fitness. To test genome replication, iSLK WT cells were depleted of PAPα/γ or ZFC3H1, and ORF57 DNA levels were monitored over time by qRT-PCR. Depletion of PAPα/γ or ZFC3H1 had no significant effect on the levels of viral genome replication at 48 hpi (Fig 4D). We next investigated whether PPD inactivation affects production of infectious virions. To test this, we collected media from iSLK WT cells at 0, 8, 12, 24, 48 and 72 hours after lytic reactivation and used it to infect HEK293 cells. Two days later, viral infection was analyzed by flow cytometry to detect the GFP expressed by BAC16. Most of the HEK293 cells infected with media collected from iSLK WT cells at 48 and 72 hpi were GFP positive (Fig 4E). Importantly, depletion of PAPα/γ or ZFC3H1 had no effect in the production of infectious virus as the percentage of GFP positive cells was similar to that of cells treated with a control siRNA (Fig 4E). These data suggest that the premature expression of late genes observed upon PPD inactivation does not dramatically perturb the virus life cycle in cultured cells.

### NRDE2 is needed for proper expression of late genes at 48 hours post lytic reactivation

To determine how late transcripts avoid degradation at their proper time of expression, we centered our attention on the human <u>n</u>uclear <u>R</u>NAi-<u>de</u>fective <u>2</u> (NRDE2) protein. NRDE2 localizes to nuclear speckles where it forms a 1:1 complex with MTR4 to inhibit its recruitment and RNA degradation (54). Given this protective activity, we hypothesized that KSHV uses NRDE2 to protect late transcripts from degradation. Therefore, we depleted cells of PAPα/γ, NRDE2 (Fig 5A) or both simultaneously and measured expression levels of late transcripts by qRT-PCR at 24 hpi (Fig 5B) or 48 hpi (Fig 5C). As expected, PAPα/γ depletion resulted in increased expression levels of late genes at 24 hpi but no effect at 48 hpi (Fig 5B and C) (green bars). In contrast, NRDE2 depletion caused a reduction in expression levels of all late transcripts at 48 hpi but had no effect at 24 hpi (Fig 5B and C) (orange bars) suggesting that NRDE2 protects KSHV late transcripts from PPD-mediated degradation at 48 hpi. Importantly, co-depletion of PAPα/γ and NRDE2 restored late transcripts levels to that of control siRNA treated cells (Fig 5C) (blue bars). We conclude that after viral genome replication, the host NRDE2 protects KSHV RNAs from PPD.

**Fig 5.**
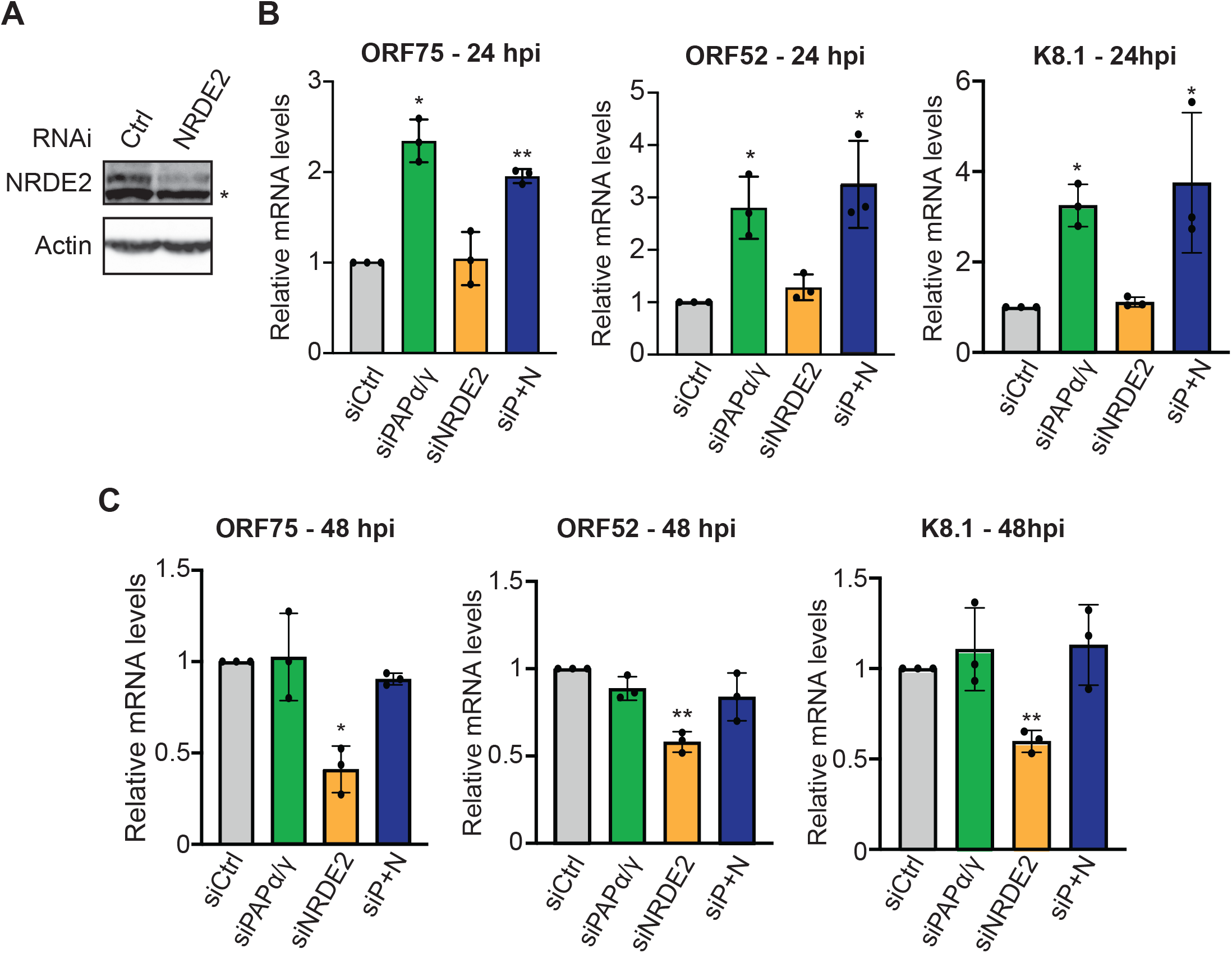
NRDE2 protects viral late transcripts from degradation. (A) NRDE2 knockdown efficiency was determined by quantitative western blot. Actin serves as loading control. (*) nonspecific band. (B and C) Bar graphs showing relative ORF75, ORF52 and K8.1 mRNA levels at 24 (B) and 48 (C) hpi in iSLK WT depleted of PAPα/γ (green), NRDE2 (orange) and PAPα/γ and NRDE2 combined (blue). Total RNA was harvested 24 or 48 hpi and analyzed by qRT-PCR. Values are displayed relative to siCtrl after normalization to the 18S rRNA level. All values are averages, and the error bars are standard deviations (n = 3). *P* values were determined by two-tailed unpaired Student’s *t* test: * < 0.05; ** < 0.01; *** < 0.001.

## Discussion

KSHV transcripts are subject to degradation by at least two host-mediated nuclear RNA decay pathways, PPD and an ARS2-dependent decay pathway (29). KSHV ORF57 increases viral transcript stability by protecting RNAs from ARS2-dependent decay (29). Our work here suggests that KSHV uses PPD to post-transcriptionally control the premature accumulation of late transcripts during the early stages of the viral lytic phase. In the context of PPD inactivation, late transcripts aberrantly accumulate at 24 hpi (Fig 1 and 2), but the premature production of late transcripts does not require functional ORF24. Therefore, the transcripts do not accumulate as a secondary consequence of the up-regulation of the late gene inducer ORF24 (Fig 3). Presumably, the open chromatin and high transcription of the viral genome during early lytic phase allows low-level cryptic transcription of late genes (Fig 6A). The transcripts are eliminated by PPD, so no proteins are produced. At their proper time of expression, late transcripts evade PPD by an NRDE2-dependent mechanism (Fig 5). Thus, we propose that KSHV fine-tunes temporal expression of its genes using a specific host RNA quality control pathway.

**Fig 6.**
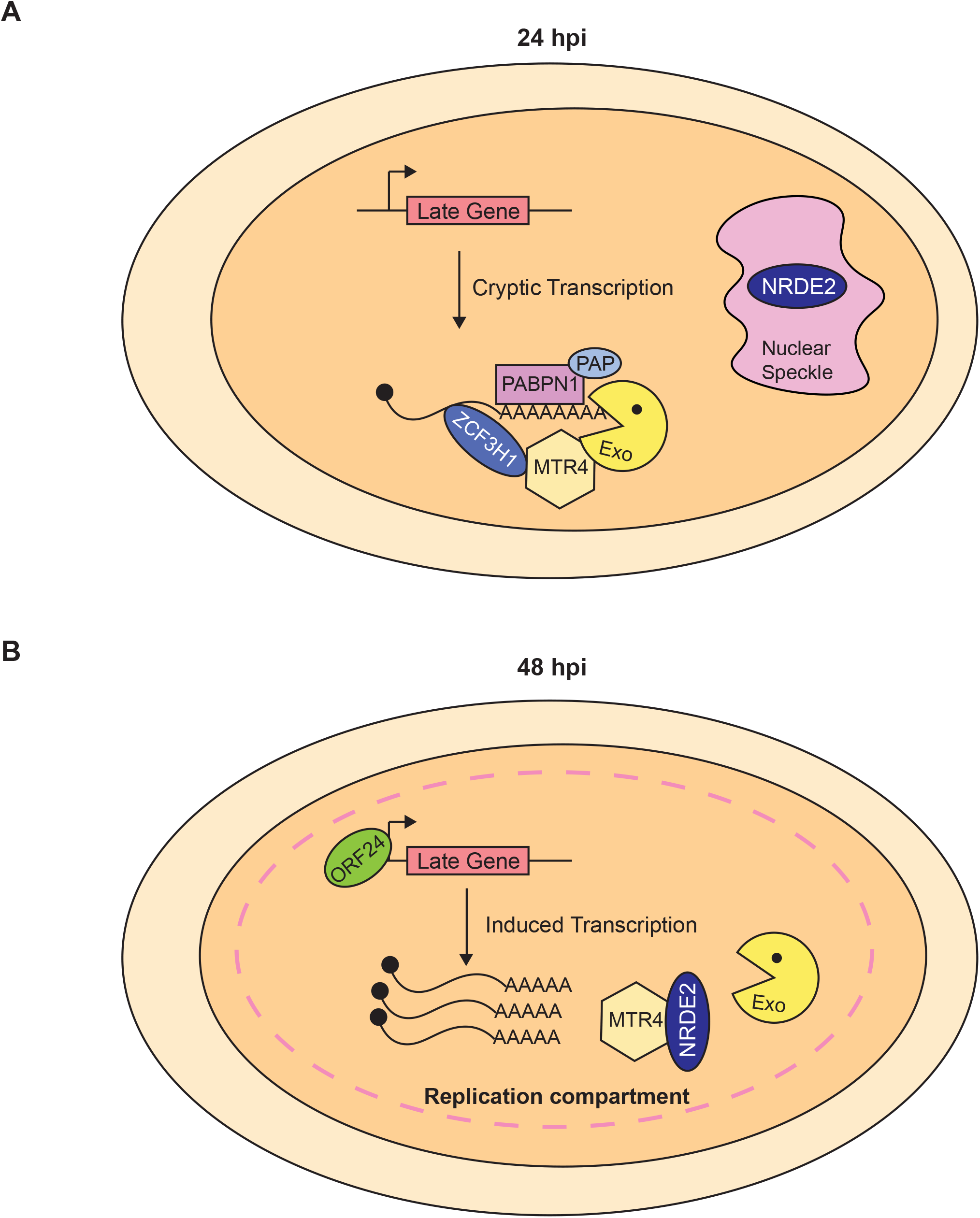
Model of PPD regulation of KSHV late genes. (A) KSHV late genes are cryptically transcribed at 24 hpi, but the transcripts do not accumulate due to PPD. (B) At 48 hpi, KSHV late transcripts evade PPD and accumulate at high levels. We speculate that KSHV replication compartments coalesce with nuclear speckles where NRDE2 protects viral transcripts from PPD degradation by sequestering MTR4.

Increasing evidence shows that RNA decay pathways play a critical role in controlling viral infection (29, 55–57). The nuclear RNA decay factors, MTR4 and ZCCHC7, translocate to the cytoplasm where they promote exosome-mediated degradation of viral transcripts of multiple RNA virus (55). In the case of KSHV, the ORF57 protein protects viral transcript from ARS2-dependent decay by preventing MTR4 recruitment (29). In these examples, the viruses must circumvent the RNA QC machinery to properly express its genes. Here we propose that the virus hijacks PPD activity to fine-tune the temporal expression of its genes. That is, KSHV allows PPD to degrade late transcripts that arise as a consequence of cryptic transcription at 24 hpi (Fig 2 and 6A). However, at the appropriate time of expression, KSHV blocks PPD so late genes are expressed (Fig 4A-C). The long co-evolution of herpesviruses with their specific hosts has selected for sophisticated host-pathogen interactions. These contrasting interactions with the host RNA QC pathways represent intriguing examples of virus-host co-evolution that ensures KSHV expresses its genes in a precise temporal manner.

Our observations also contribute to the understanding of the distinctions between host PPD, PAXT and ARS2-mediated decay processes. While it seems likely that PPD and PAXT represent the same pathway, it has been reported that ARS2 is involved in PAXT recruitment to unstable transcripts (20). However, in the context of KSHV infection, PPD and an ARS2-mediated decay pathway appear to be independent pathways. For example, simultaneous depletion of PAPα/γ and ARS2 resulted in greater stabilization of viral transcripts than depletion of either alone during iSLK-ΔORF57 reactivation (29). Furthermore, we show that depletion of PAPα/γ enhances specific viral genes, while ARS2 depletion has little effect (Figs 1 and 2). Importantly, we show that depletion of ZFC3H1, a PAXT component, mimics PAPα/γ depletion as the same group of viral genes are upregulated (Fig 2). Overall, these data suggest that PPD and PAXT represent the same process, but ARS2 is not absolutely required for PPD/PAXT-mediated decay. However, more work is needed to completely define the overlap and independence of these RNA QC pathways.

PPD inactivation results in the aberrant temporal expression of KSHV late genes. However, this atypical expression of late genes does not affect viral genome replication or production of infectious virions in our iSLK cells (Fig 4D & E). Several reasons may explain this unexpected result. Even though PAP knockdown increases late gene expression levels relative to cells treated with a control siRNA, the expression levels reached may not be high enough to affect viral fitness. Indeed, expression of late genes at 48 or 72 hpi is considerably higher than at 24 hpi (Fig 4A-C). Consequently, the levels of late genes reached at 24 hpi may not be sufficient to disrupt viral physiology in iSLK cells. Another possibility is that the virus uses PPD to keep expression levels of late genes low during early phases of lytic infection as these may elicit an immune response in an infected organism. Indeed, circulating anti-K8.1 antibodies detected in Kaposi’s sarcoma patients support that this PPD-restricted gene elicits an immune response (58–60). If the role of PPD is to keep potentially immunogenic genes low, we would miss this phenotype in a cell culture system.

Late transcripts evade PPD during late phases of replication, but the mechanism of PPD evasion is not completely understood. Our data show that the host factor NRDE2 is required for evasion of PPD during late infection (Fig 5). NRDE2 function is linked to its residence in nuclear speckles, where it interacts with MTR4 to inhibit its activity thereby protecting speckle-associated mRNAs (54). As expression of KSHV late genes occurs at the onset of viral genome replication, we speculate that NRDE2 re-localizes to replication compartments (Fig 6B). When this occurs, NRDE2 interacts with MTR4 preventing the recruitment of the nuclear exosome to viral transcripts. Consequently, late transcripts evade PPD and are properly expressed. Supporting this idea, HSV1 replication compartments coalesce with nuclear speckles (61), but whether this occurs in KSHV has yet to be tested.

Additional factors may contribute to protection of late transcripts from PPD during late phases of KSHV reactivation. For example, expression of KSHV late genes requires the action of several viral transactivation factors (48–52). ORF24 is essential to recruit RNA pol II and other viral transactivation factors to late gene promoters (51). In principle, ORF24-induced transcription may promote the co-transcriptional recruitment of factors that protect transcripts from degradation. Evasion and exploitation of nuclear RNA decay pathways by KSHV is only beginning to be understood. Further experimentation is needed to substantiate the role of NRDE2 and identify other factors involved in these processes.

## Materials and Methods

### RNA-seq: library preparation

iSLK WT cells were transfected with a non-targeting control siRNA or a two-siRNA pool targeting PAPα and PAPγ (PAPα/γ), ARS2 or both PAPα/γ and ARS2 combined (dKD) using the concentrations specified in the siRNA transfection section. Total RNA was harvested three days after siRNA transfection and 24 hours post lytic reactivation. One μg of intact total RNA per condition was used to make stranded mRNA-seq libraries with the Stranded mRNA-Seq kit (KAPA Biosystems) as per manufacturer’s protocol. The strand-specific single-end RNA-sequencing was performed using Illumina HiSeq2500.

### RNA-seq analysis

The qualities of sequencing reads were evaluated using NGS QC Toolkit (v2.3.3) (62) and high-quality reads were extracted. The human reference genome sequence and gene annotation data, hg19, were downloaded from Illumina iGenomes (https://support.illumina.com/sequencing/sequencing_software/igenome.html). The viral genome was downloaded from NCBI GenBank (https://www.ncbi.nlm.nih.gov/nuccore/GQ994935.1). The qualities of RNA-sequencing libraries were estimated by mapping the reads onto human transcript and ribosomal RNA sequences (Ensembl release 89) using Bowtie (v2.2.9) (63). STAR (v2.5.2b) (53) was employed to align the reads onto the human and viral genomes, Picard (v1.140) (https://broadinstitute.github.io/picard/) was employed to sort the alignments, and HTSeq Python package (64) was employed to count reverse-stranded reads per gene. DESeq2 R Bioconductor package (65) was used to normalize read counts and identify differentially expressed (DE) genes. The resulting gene expression analyses are given in Supplementary Tables S1 for viral genes exclusively and S2 for human plus viral genes. The enrichment of DE genes to pathways and GOs were calculated by Fisher’s exact test in R statistical package. Genome coverages were calculated using SAMtools (v0.1.19) (66), BEDTools (v2.26) (67), and bedGraphToBigWig (https://genome.ucsc.edu/index.html). The heatmap was generated using Morpheus from the Broad Institute (https://software.broadinstitute.org/morpheus/).

The data discussed in this publication have been deposited in NCBI’s Gene Expression Omnibus (68) and are accessible through GEO Series accession number GSE144747 (https://www.ncbi.nlm.nih.gov/geo/query/acc.cgi?acc=GSE144747).

### Cell Culture

iSLK cells were grown at 37°C with 5% CO_2_ in DMEM (Sigma) supplemented with 10% Tet-Free fetal bovine serum (FBS, Atlanta Biologicals), 1x penicillin-streptomycin (Sigma), and 2 mM L-glutamine (Fisher). iSLK WT cells were grown in the presence of 0.1 mg/mL G418 (Fisher), 1 µg/mL puromycin (Sigma) and 50 µg/mL hygromycin. iSLK-ORF24^R328A^ cells (gift from Dr. Britt Glaunsinger, University of California Berkeley) were grown under the same conditions, except 200 µg/mL of hygromycin was used.

### siRNA Transfections

iSLK cells were transfected with 20 or 40 nM siRNA (Silencer Select, ThermoFisher) using RNAiMAX transfection reagent (Invitrogen) per manufacturer’s instruction. Specifically, we used final concentrations of 40 nM siRNAs for ZFC3H1, MTR4 and NRDE2 and 20 nM siRNAs ARS2. For PAPα/γ, we used 20 nM each of siRNAs that target PAPα or PAPγ for a total of 40 nM siRNA. Twenty-four hours after siRNA transfection, cells were split into new plates and allowed to grow for another 24 hours, after which doxycycline and NaB was added to induce lytic reactivation. Thus, total RNA was harvested 72 hours post siRNA transfection and 24 hours post lytic reactivation. Nontargeting control, PAPα/γ, ZC3H1 and MTR4 siRNAs are the same as previously used (29). NRDE2 siRNAs are: 5’ GGUGUUGUUUGAUGAUAUUtt 3’ (s30063) and 5’ GUUUAGUACCUUUUCGAUAtt 3’ (s30064).

### Quantitative RT-PCR

RNA was harvested using TRI reagent (Molecular Research Center, Inc.) according to the manufacturer’s protocol. Following extraction, RNA was treated with RQ1 DNase (Promega). Oligo dT_20_ was used to prime cDNA synthesis with MuLV reverse transcriptase (New England Biolabs). Real-time reactions used iTaq Universal SYBR Green Supermix (Biorad). Primers are listed in Table S3.

### KSHV Reactivation and Infection

Lytic reactivation of iSLK derived cells was achieved by adding doxycycline (1 µg/ml) and NaB (1mM). Tissue culture supernatants from iSLK WT cells were collected at 0, 8, 12, 24, 48 and 72 hpi, centrifuged for 5 min at 1000 × g and passed through a 0.45 um filter. Polybrene was added (8 µg/mL final concentration), and 300 µL were applied to HEK293 cells grown in a 12-well plate. Cells were centrifuged for 45 min at 30°C and then incubated in 5% CO_2_ at 37°C for 2 hours. After this, media was replaced and cells were analyzed by flow cytometry 24 hours later.

### Western Blotting

Cells were lysed in buffer containing 100 mM NaCl, 50 mM Tris-HCl pH 7.4, 1% Triton X-100, 1X Protease Inhibitor cocktail (PIC) (Calbiochem) and 250 μM PMSF. Proteins were resolved by SDS-PAGE and analyzed by western blot using standard procedures. Antibodies used are rabbit polyclonal anti-ARS2 (Abcam, ab192999), rabbit polyclonal anti-MTR4 (Abcam, Ab70551), rabbit polyclonal anti-NRDE2 (Proteintech, 24968) and mouse monoclonal anti-Actin (Abcam, ab6276). Quantitative westerns were performed using infrared detection with an Odyssey Fc and quantification was performed using ImageStudio software (LI-COR Biosciences).

## Acknowledgements

We thank Anna Scarborough and Juliana Flaherty for critical review of this manuscript. We thank Dr. Divya Nandakumar, Jennifer Blancas and Dr. Britt Glaunsinger (UC Berkeley) for the iSLK ORF24^R328A^ cells. We thank Spencer Barnes (UTSW) for GEO submission. The work was supported by NIH/NIAID R01 AI123165 (to N.K.C.), the Welch Foundation I-1915-20170325 (to N.K.C) and Cancer Prevention and Research Institute of Texas RP150596 (to the Bioinformatics Core Facility, UTSW).

---

**Table S1. Differential expression of KSHV genes.** This spreadsheet contains the expression levels of all KSHV genes in samples depleted of PAPα/γ, ARS2 and dKD relative to siCtrl.

**Table S2. Differential expression of human and KSHV genes.** This spreadsheet contains the expression levels of human and KSHV genes in samples depleted of PAPα/γ, ARS2 and dKD relative to siCtrl.

**Table S3. Primers used in this study.** Target, sequence, and primer number (ID) for all PCR primers used herein.

